# Oxidative stress changes interactions between two bacterial species

**DOI:** 10.1101/2023.05.24.542164

**Authors:** Rita Di Martino, Aurore Picot, Sara Mitri

## Abstract

Knowing how species interact within microbial communities is crucial to predicting and controlling community dynamics, but interactions can depend on environmental conditions. The stress-gradient hypothesis (SGH) predicts that species are more likely to facilitate each other in harsher environments. Even if the SGH gives some intuition, quantitative modeling of the context-dependency of interactions requires understanding the mechanisms behind the SGH. In this study, we show with both experiments and a theoretical analysis, that varying the concentration of a single compound, linoleic acid, modifies the interaction between two bacterial species from competitive at a low concentration, to facilitative at higher concentrations where linoleic acid becomes toxic for one of the two species. We demonstrate that the mechanism behind facilitation is that one species is able to reduce Reactive Oxygen Species (ROS) that are produced spontaneously at higher concentrations of linoleic acid, allowing for short-term rescue of the species that is sensitive to ROS and longer coexistence in serial transfers. In our system, competition and facilitation between species can occur simultaneously, and changing the concentration of a single compound can alter the balance between the two.

## Introduction

Multispecies microbial communities colonize almost every environment. Despite their small individual size, microbes can form very large populations that significantly affect their surroundings. For example, they can greatly influence the health and behavior of their living hosts (1) – for better or worse – or alter the physical and chemical properties of surfaces they are living on (2). How these effects play out depends on a community’s species composition and how it changes over time, which in turn depends on how the different species interact: who drives whom extinct and who promotes whose growth. But how we expect microbial species to interact with one another, what drives their interactions, and how interactions shape long-term coexistence remain matters of debate (3–10).

Here, we define interactions between species as the effect of one species on the growth and death of another and focus on environmentally-mediated interactions (11). A species may negatively affect another by making the environment worse for it, for example by depleting a common resource, or releasing a toxic waste product. Alternatively, a species may affect another positively by improving the environment for the other, or for itself, accidentally benefiting the other (11, 12). Such positive effects can happen through detoxification, cross-feeding, or the release of public goods such as siderophores (13).

Whether or not a species improves the environment or makes it more difficult for another species to grow will depend on the properties of the environment itself, suggesting that interactions should be context-dependent (12, 14) – assuming that one can correctly discern true context-dependency from background confounding effects (15). Indeed, several studies have demonstrated that changes to the chemical composition of the environment can alter interaction sign (16–23).

Ultimately, our understanding of interspecies interactions should allow us to predict the long-term composition of microbial communities, i.e. which species are likely to coexist. Coexistence theory usually considers competitive interactions, although recent approaches also integrate facilitation and mutualism (24). In particular, Modern Coexistence Theory (25, 26) focuses on species-species interactions with a general Lotka-Volterra framework, while Contemporary Niche Theory (27, 28) sees interspecific interactions as mediated by an environment from a consumer-resource perspective (29), which takes the dependency of interactions on environmental conditions into account (11, 30). Indeed, coexistence outcomes can critically depend on the mechanistic details of interactions (24, 31, 32).

One example where context-dependency has been proposed is the Stress-Gradient Hypothesis (SGH), which states that positive interactions between species are likely to increase with environmental stress. The SGH was originally described in plants (33, 34), but has also been observed in bacterial communities (16, 35–37). In our previous work (36), we measured pairwise interactions between four bacterial species that were isolated from polluting industrial liquids called Metalworking Fluids (MWF). We showed that all pairwise interactions were positive in toxic MWF, but became more competitive when the environment was made more benign. Although this result supports the SGH, the complexity of the MWF medium and its unknown chemical composition prevented us from identifying a mechanism to explain facilitation in this toxic environment.

In this study, we tested the SGH in a simpler system, combining experiments and mathematical models to study species interactions and coexistence in environmentally defined conditions. Our system consists of two bacterial species originally isolated from MWF, *Agrobacterium tumefaciens* (henceforth *At*) and *Comamonas testosteroni* (*Ct*), growing in a defined medium containing linoleic acid (LA), a fatty acid commonly present in many MWFs (38) as the sole carbon source. We chose this carbon source as it becomes toxic for *At* – but not *Ct* – at high concentration, making it suitable to explore the relationship between toxicity and interactions, and to mechanistically test the SGH and how facilitation affects coexistence.

We found that the toxicity of LA for *At* was due to the accumulation of Reactive Oxygen Species (ROS). Using the models, we show how to drive the interactions in the two-species co-culture towards more competition or facilitation by simply reducing or increasing the initial LA concentration, respectively, in line with SGH. Moreover, by artificially reducing environmental toxicity (by adding an antioxidant) at high LA concentration, we were also able to revert back to competition, hence mechanistically showing how the SGH works in our system. We then use our model to predict how interactions would affect long-term coexistence and found that toxicity extends the duration of coexistence in the short term, which was experimentally validated. Overall, the simplicity of our system allowed us to identify the mechanism behind the interactions between our bacteria and to shape them just by manipulating toxicity through the concentration of a single chemical compound.

## Results

### Linoleic acid has concentration-dependent effects in monoculture

To build on our previous work (36) and track how pair-wise interactions change with varying compound concentration and varying toxicity, we designed a simpler medium containing a single MWF compound at a time. We selected ten compounds commonly found in MWF and tested their effect on *At* and Ct in monoculture at different concentrations (Fig. 1A). As *Ct* was facilitating *At* in MWF (36), we were looking for compounds on which *Ct* could grow but that would be challenging for *At*. Linoleic acid (LA) was a good candidate, as it acted as a nutrient source for both species at low concentration, while at high concentration, it was toxic for *At* even though *Ct* could still grow well (Fig. 1B). To generate quantitative predictions on the behavior of these two species in mono- and co-culture, we developed a mathematical model (Ordinary Differential Equations (ODEs) describing dynamics of species and substrate abundances, see Methods) with parameters (e.g. growth and death rates) that best fit the mono-culture growth curves (Fig. 1B). To capture the response of *At* to LA in the model, LA acted as a nutrient source that could also cause death at increasing concentrations (see Methods). We then used the model to predict how the two species are expected to interact if co-cultured at different LA concentrations (Fig. 2A, B).

**Fig. 1.**
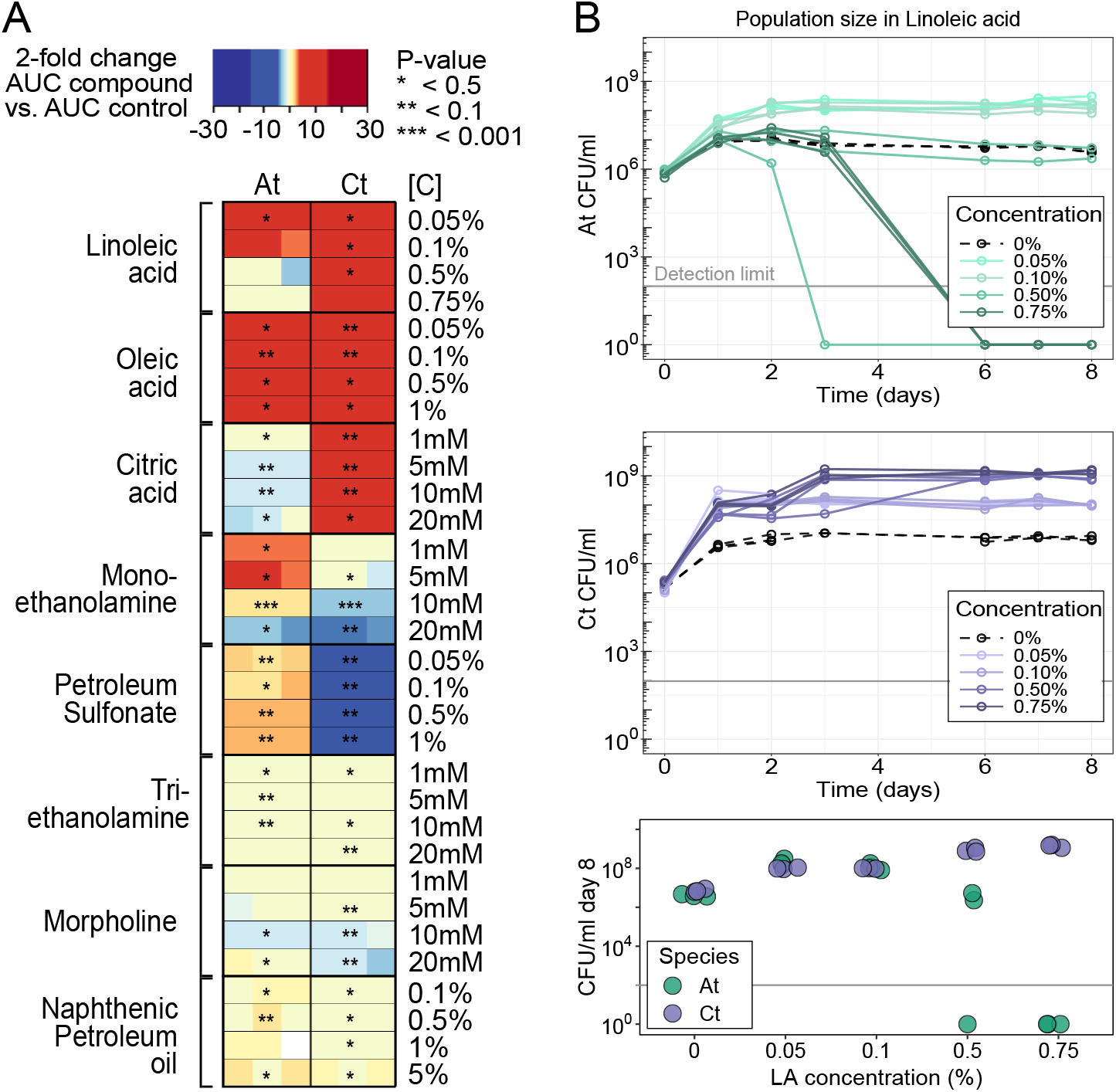
(A) Growth of *A. tumefaciens* (*At*) and *C. testosteroni* (*Ct*) in presence of different compounds at a range of concentrations in triplicates (each replicate is one colored square in the rectangle). The tested compound is the only carbon source in the medium (see Methods). Heatmaps show the fold change between the area under the growth curve (AUC) of each sample replicate and the AUC of the mean of the control replicates where no compound was added. Blue shades represent negative fold change (bacteria grew to significantly smaller populations than the control) and orange shades represent positive fold change (bacteria grew to greater population sizes than the control). Statistical significance is based on t-tests (*: *P <* 0.05, **: *P <* 0.01, ***: *P <* 0.001). (B) Growth curves that generated the linoleic acid data in panel A. We observe some growth for both species in the absence of any carbon source (0%). While we were unable to find the source of this growth, similar observations have been previously reported (39–41). All three technical replicates per condition are shown. The bottom panel shows population sizes at day 8 for a clearer comparison.

**Fig. 2.**
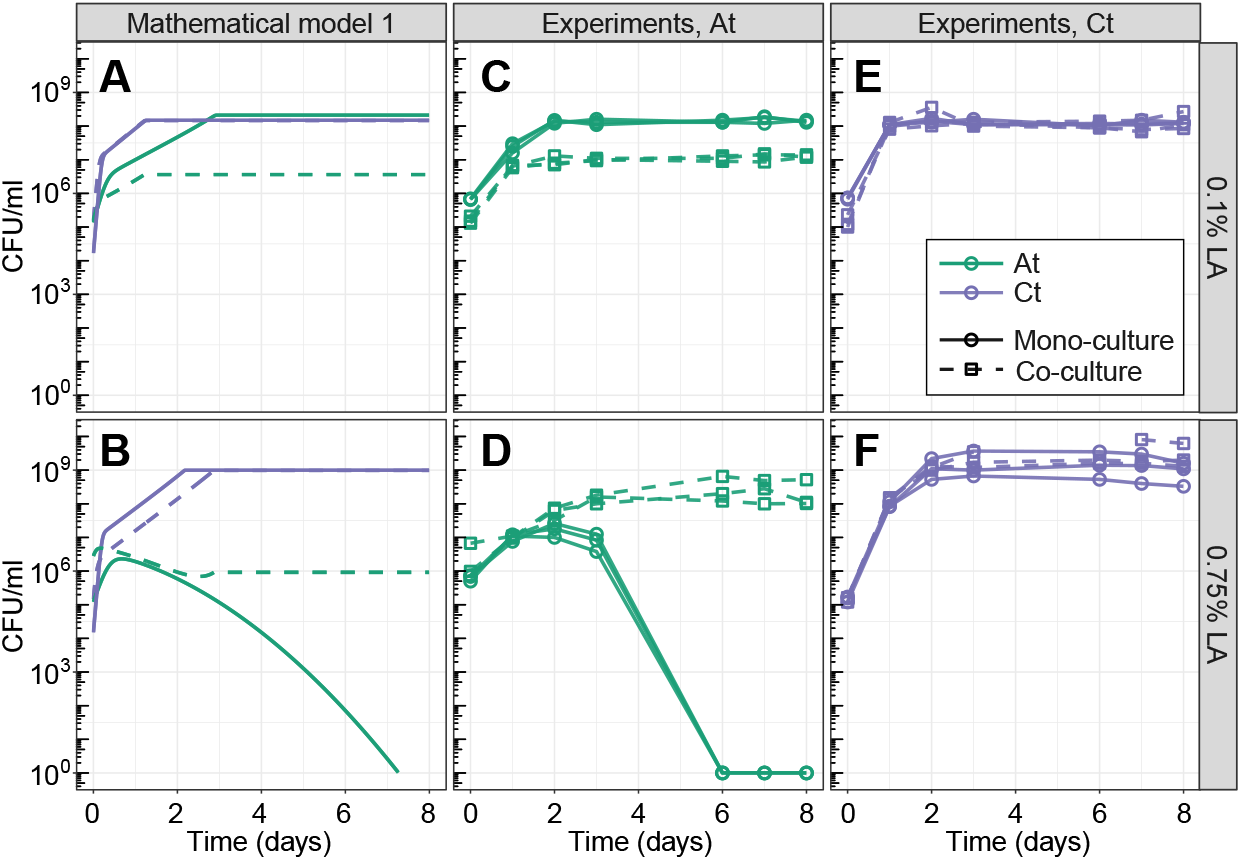
Growth of the two species in mono- and co-cultures at the two LA concentrations, predicted by the model and in experiments. (A, B) Predictions of model 1, where LA is a nutrient but becomes increasingly toxic over time. Model parameters were estimated by fitting to the mono-culture data (see Methods). (C-F) Experiments showing the growth of *At* (C, D) and *Ct* (E, F) at 0.1% LA (C, E) or 0.75% LA (D, F). Both species survive in monoculture at 0.1% LA, whereas *At* grows and then dies in mono-culture at 0.75%. At 0.1% LA, *At* suffers from the presence of *Ct* in co-culture, while *Ct* ‘s growth is not significantly affected. At 0.75% LA, *At* is rescued by Ct in the co-culture, and *Ct* ‘s growth is not significantly affected. The model does a reasonable job at capturing the overall dynamics, but underestimates *At* ‘s growth in co-culture at 0.75% LA. For experimental data, all three technical replicates per condition are shown. See main text for statistics.

### Linoleic acid concentration determines interaction sign

The model predicted that increasing the concentration of LA in a co-culture of the two species could change the interaction sign from negative to positive. More specifically, at low concentration, both species compete for the sole nutrient source LA. Once the concentration becomes high enough to kill *At*, however, we expect to observe facilitation, as *Ct* consumes LA and reduces its concentration, making the environment less toxic for *At* (Fig. 2A, B). We tested this prediction in the lab by growing *At* and *Ct* in mono- and co-culture in 0.1% and 0.75% LA as low and high LA concentrations, respectively. The results were in line with the predictions of the model: at low LA concentration (0.1%), *At* grew significantly to smaller population sizes in the co-culture compared to mono-culture (AUC of *At* at 0.1% LA in monoculture: 9× 10^8^ *±*4 ×10^7^ vs. co-culture: 7 ×10^7^*±* 9 ×10^6^, t-test *P <* 0.001, Fig. 2C), showing that there was competition for LA. At high LA concentration (0.75%), the presence of *Ct* in the co-culture rescued *At*, allowing it to grow to population sizes that were orders of magnitude greater than alone (AUC of *At* at 0.75% LA in mono-culture: 4 ×10^7^ *±*2 ×10^7^ vs. co-culture: 2× 10^9^*±* 10^9^, t-test *P* = 0.03, Fig. 2D). The growth of *Ct* was not significantly affected by *At* in either condition (AUC of *Ct* in 0.1% LA in mono-culture: 9 ×10^8^*±* 5 ×10^7^ vs. co-culture: 9 ×10^8^ *±*10^8^, t-test *P* = 0.98; AUC of *Ct* in 0.75% LA in mono-culture: 10^10^ *±*8 ×10^9^ vs. co-culture: 2 ×10^10^*±* 2 ×10^10^, t-test *P* = 0.42, Fig. 2E, F).

Although these results matched the model predictions qualitatively, *At*’s growth in co-culture was greatly underestimated by the model (Fig. 2B, D, green dashed lines). Furthermore, using the parameters estimated from all mono-cultures, the model does not correctly predict the hump-shaped growth of *At* at 0.75% LA (Fig. 2D), even though it already assumes the accumulation of toxicity. This suggests that estimating model parameters where both growth and death are caused by a single compound is challenging. We next focused on exploring the mechanism behind this possible increase in toxicity, in particular if we can decouple toxicity from the linoleic acid itself.

### ROS accumulates upon oxidation of linoleic acid and causes death of *At* unless *Ct* is present

Based on the literature, we hypothesized that spontaneous oxidation of LA might release reactive oxygen species (ROS) (42–45). Accordingly, we used the Thiobarbituric Acid Reactive Substances (TBARS) assay (see Methods) to test for the presence of ROS in our system (46). Indeed, at both LA concentrations of the cell-free medium, ROS accumulated over time (Fig. 3B, cell-free), supporting the idea of a chemical reaction leading to ROS production, such as spontaneous oxidation due to exposure of light and air (43, 45). ROS were significantly less abundant in *Ct* monoculture and in the co-culture than in the monoculture of *At* (*Ct* mono-culture vs. *At* monoculture, Tukey’s test for multiple comparisons *P* = 0.0002; *Ct* co-culture vs. *At* mono-culture, *P* = 0.0002, Fig. 3B). The lack of ROS accumulation whenever *Ct* was present led us to hypothesize that *Ct* neutralizes ROS, reducing environmental toxicity and rescuing *At* in co-culture, allowing it to survive and grow (Fig. 3A shows an experimental repeat of 2D, F). These results also showed that toxicity was not caused by the increase of LA concentration itself, but rather by the accumulation of ROS. We adapted our model accordingly, leaving LA as a nutrient exclusively and adding ROS as an additional toxic compound that is generated through spontaneous LA oxidation. The co-culture growth prediction is qualitatively captured in this updated version of the model, in particular the growth of At in high LA concentration is not under-predicted as it was in model 1 (Fig. 3A compared to Fig. 2B).

**Fig. 3.**
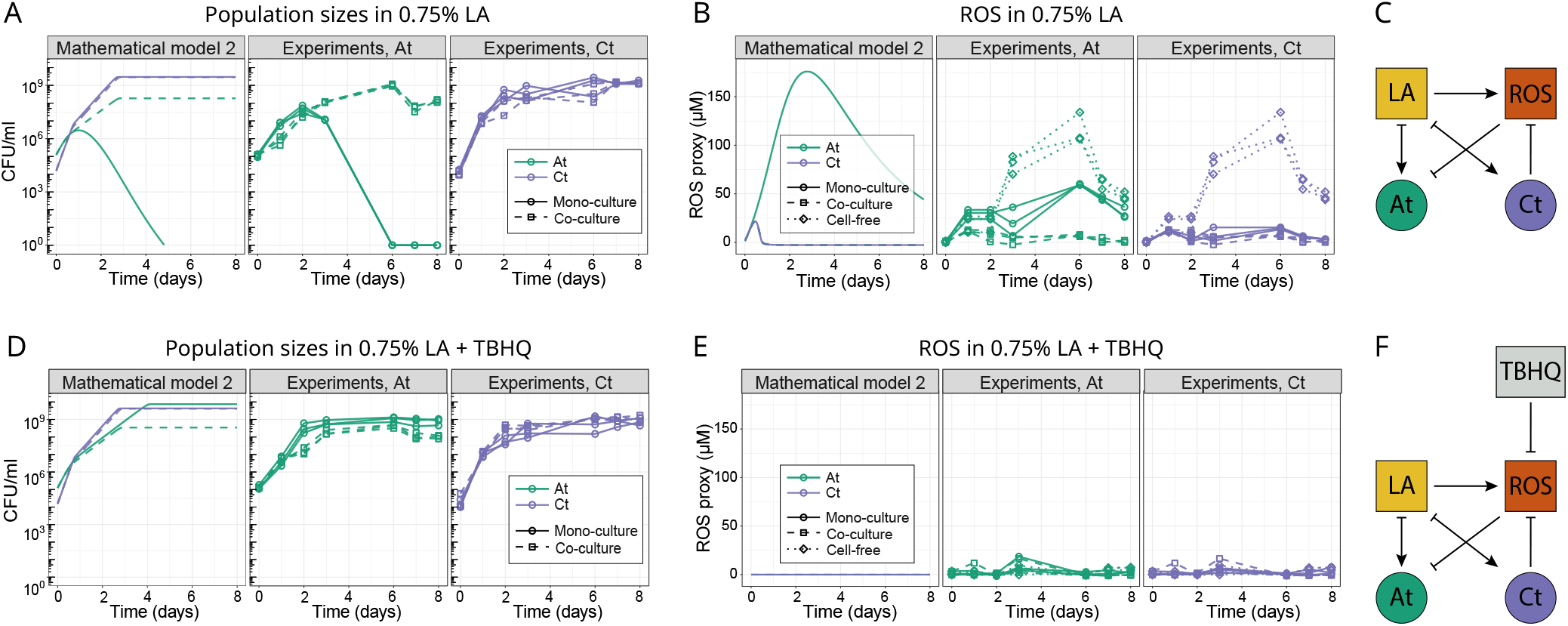
Comparing population and ROS abundance over time. (A, D) Population sizes of both species in mono- and co-culture at 0.75% LA (A) or 0.75% LA with the antioxidant TBHQ added daily (D) (see Methods). Panel (A) shows a biological repeat of the experiment shown in Fig. 2, but where ROS was measured simultaneously (panel B). The mathematical model in the left panels of (A) and (D) is the second implementation of the model, where ROS is explicitly modelled as a separate chemical from LA. In model 2, LA is only a resource (see Methods). Model parameters were estimated by fitting to the mono-culture data (see Methods). (B, E) A proxy for ROS concentration (MDA-TBA2, see Methods for assay) over time in the different experimental treatments and in model 2 (left panel) in the 0.75% LA medium (B) or where TBHQ was added (E). The cell-free and co-culture data are identical in the *At* and *Ct* subpanels, as they come from the same samples. For experimental data, all three technical replicates per condition are shown. (C, F) A diagrammatic representation of the relationships between chemicals (in squares) and bacterial species (circles): (C) LA is consumed as a resource by both species, and generates ROS. ROS inhibits *At* but can be reduced by *Ct*. (F) TBHQ inhibits ROS, making ROS removal by *Ct* superfluous. ROS removal by TBHQ or by *Ct* in co-cultures, rescues *At*. This translates into facilitation of *At* by *Ct* in the top row and competition in the bottom row, where TBHQ removes ROS.

### Antioxidant rescues *At* and reverses the interaction between *At* and *Ct*

We used this second model to explore what would happen if we removed the toxicity by setting the production of ROS to 0 and leaving LA strictly as a nutrient. This led to two predictions: in absence of toxicity, (i) at 0.75% LA, *At* should survive even in monoculture and reach a higher population size compared to 0.1% LA, and (ii) we should observe competition between *At* and *Ct* even at 0.75% LA (Fig. 3D, left). To test these predictions, we first added an antioxidant molecule to the cell-free LA medium to verify whether its presence would decrease ROS concentration. We chose to use tert-butylhydroquinone (TBHQ) for its antioxidant properties (47) at a concentration that did not inhibit bacterial growth. In addition to the cell-free medium, we also added TBHQ to *At* and *Ct* monocultures, and the co-culture at the beginning of the growth assay and every 24 hours to have a regular input of fresh antioxidant, mimicking continuous ROS neutralization by *Ct*. We found that TBHQ successfully decreased ROS concentration in all tested culture conditions compared to their value in standard 0.75% LA (Fig. 3B, E). In support of prediction (i), adding TBHQ rescued *At* in monoculture (Fig. 3D, center), allowing it to reach a significantly higher population size in 0.75% LA + TBHQ compared to 0.1% (AUC of *At* mono-culture in 0.1% LA vs. 0.75% LA+TBHQ, t-test *P* = 0.013, compare solid lines in Fig. 3D, center and Fig. 2C). We suppose that in this ROS-free condition *At* could exploit the greater availability of LA as a nutrient. And as per prediction (ii), in 0.75% LA + TBHQ *At* grew to significantly lower population sizes in the presence of *Ct* than in monoculture (Fig. 3D, center, AUC of *At* mono-culture in 0.75% LA+TBHQ: 3.7 × 10^9^ *±*1.3× 10^9^ vs. co-culture: 7.9 × 10^8^*±* 1.5× 10^8^, *df* = 5, t-test *P* = 0.019), meaning that the interaction between *At* and *Ct* switched from facilitation to competition in the absence of toxicity. Overall, both model predictions were confirmed, demonstrating how we can shape the interaction between *At* and *Ct* by manipulating the level of toxicity in the environment.

### Separation of nutrient and toxic components in the model makes short-term coexistence between *At* and *Ct* more likely

In the first version of the model, toxicity depended on LA concentration, while in the second, it emerged on ROS accumulation, while LA acted exclusively as a nutrient. While this difference may seem like an implementation detail, we know from early theoretical work that one resource and one inhibitor can allow two species to coexist, while a single compound can only do so under very restrictive conditions (31, 32). We sought to verify this prediction and explore whether coexistence between *At* and *Ct* would be possible in either version of the model and in the experiments.

We first extended the models to simulate a transfer experiment, where cultures were grown in batch for 72h and then diluted 100-fold into fresh medium and regrown, and asked how long coexistence between the two species was possible. The second version of the model including ROS predicted a larger parameter range in which the two species could coexist in the short-term (Fig. 4A). We tested this prediction experimentally by transferring 1% of the *At* and *Ct* monocultures and the co-culture in respective tubes every 72 hours in both 0.1% and 0.75% LA. After five transfers, we found a significant variation for *Ct* co-culture in 0.1% LA compared to the beginning of the transfer experiment (first vs. last transfer *Ct* co-culture 0.1% LA, t-test, *P* = 0.0029), but there was no significant change for *At* in the same condition (first vs. last transfer *At* co-culture 0.1% LA, t-test, *P* = 0.30). Moreover, in 0.1% LA, competition was still evident between *At* and *Ct*, as *At* in mono-culture maintained a significantly higher population size than in the presence of *Ct* (mono-culture *At* 0.1% LA vs. co-culture *At* 0.1% LA, t-test *P <* 0.0001, Fig. 4B, left). This suggests that coexistence between *At* and *Ct* is possible in the short-term despite the presence of negative interactions, and provides further support for our updated model. In 0.75% LA, we observed the extinction of *At* mono-culture as we expected, but we found no significant variation neither for *At* nor for *Ct* in the co-culture (co-culture *At* 0.75, t-test *P* = 0.91, co-culture *Ct* 0.75, t-test *P* = 0.2, Fig. 4B, right).

**Fig. 4.**
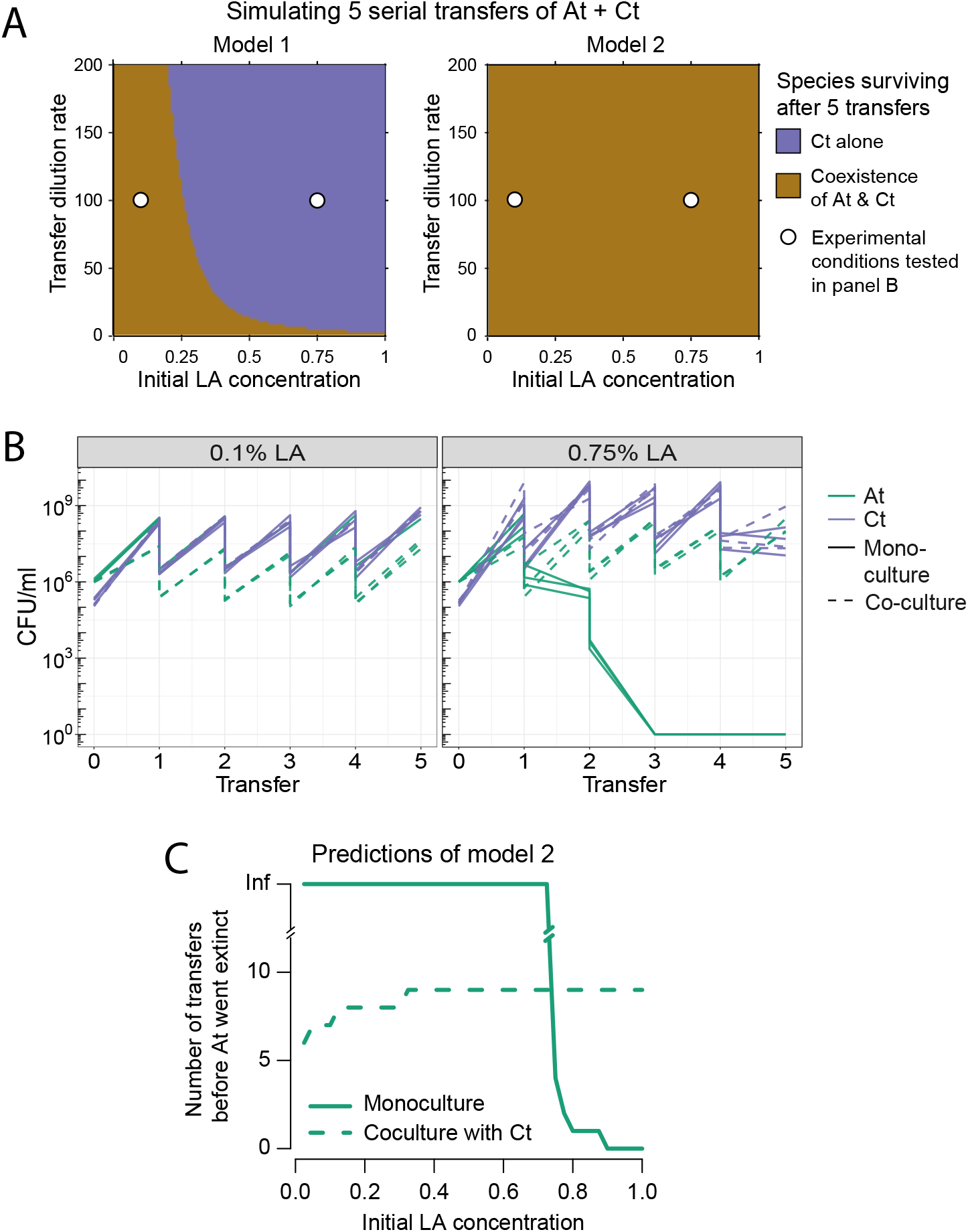
Co-existence experiments and models. (A) Prediction of short-term coexistence over 5 simulated serial transfers of the co-culture of *At* and *Ct* according to Model 1 (left) and Model 2 (right): both models allow for short-term co-existence, but the parameter space in which this is possible is larger in Model 2 (panel B, larger area representing coexistence compared to *Ct* surviving alone). White circles indicate the conditions in which experiments were run, as shown in panel B. (B) 5-Transfer experiment of *At* and *Ct* in mono- and co-culture at both 0.1% (left) and 0.75% LA (right). We show the population size in the initial culture (“transfer” 0), which is then quantified at each transfer (every 72h). We illustrate the 100-fold dilution at each transfer, although this is not explicitly quantified. All three technical replicates per condition are shown. *At* monoculture goes extinct as expected in 0.75% LA, but the two species coexist at both LA concentrations, as correctly predicted by Model 2 in panel A. (C) Model 2 predicts that *At* should survive indefinitely in mono-culture at a 100-fold dilution rate up to 0.75% LA. In co-culture, *Ct* excludes it after a few transfers at low LA concentrations, but as the concentration increases, *At* can survive for longer in cocompared to mono-culture, meaning that *Ct* facilitates *At* ‘s survival by extending its duration.

### The Stress-Gradient Hypothesis holds in simulations predicting long-term dynamics

Although the time-scale of our experiments only allowed us to explore co-existence in the short-term, it is still important to understand whether the two species would be able to coexist in the long-term. We use our final model to explore this by simulating the outcome of mono- and co-cultures of *At* and *Ct* in a gradient of initial LA concentrations. We found no conditions where the long-term stable coexistence of the two species was possible. Nevertheless, to understand whether *Ct* had an effect on the extinction time of *At*, we measured the time (transfer) at which *At* went extinct when alone and compared it to its extinction time when in co-culture with *Ct. At* lower initial LA concentrations, *At* survives indefinitely in the long-term in monoculture (Fig. 4C). As it eventually goes extinct in co-culture at all concentrations, the absence of coexistence at low concentration must be due to competitive exclusion by *Ct* (Fig. 4C). In contrast, at higher initial LA concentrations, even though *At* quickly goes extinct in mono-culture, it survives for longer in the presence of *Ct*, meaning that *Ct* facilitates *At*’s survival. We then recapitulate the SGH in the long term: *Ct* competitively excludes *At* at low LA concentrations, but facilitates it by lengthening its survival at high LA concentrations.

## Discussion

Most current research on interactions between microbes has explored positive interactions through cross-feeding (5, 10, 13, 48). Our study instead focuses on positive interactions through detoxification (11, 36), and shows how they can occur mechanistically. Facilitation through detoxification relates to the SGH. Here, with the help of our mathematical model, we show how the SGH can predict population dynamics: one species can provide a benefit to another by removing the stress that would otherwise drive the second species extinct, i.e. “niche facilitation” (24). Experimentally removing the stress from the co-culture of the two species eliminates facilitation and restores competition (Fig. 3D). This mechanistic understanding of the SGH increases its potential as a guiding principle for interspecies interactions (33, 35).

How common is this phenomenon and is it specific to the compound we focused on, linoleic acid? Fig. 1A shows that different compounds can be toxic in a species- and concentration-dependent way, suggesting that there is nothing particular about the compound we chose to study. We also know that all aerobic microbes suffer from oxidative stress to some extent, as ROS is generated as a metabolic byproduct (49), and many frequently encounter ROS in their natural habitats (50). In fact, ROS accumulation is a main plant stress response upon exposure to pathogens (51, 52), and both organisms used in this study have been associated with plants (53, 54). One reason why the role of stress in mediating microbial interactions may not receive much attention, is that lab studies are typically designed to optimize bacterial growth, which a natural environment does not.

An important gap in our study is what type of ROS is accumulating, and how *Ct* is reducing the concentration of our ROS proxy: is ROS being taken up and neutralized intracellularly or is *Ct* secreting extracellular enzymes that eliminate ROS? Answering this question is not as simple as analysing *Ct*’s genome, since all aerobic bacteria are capable of dealing with oxidative stress and we find relevant enzymes in both *Ct* and *At* (55) (Table S1). What we do know is that adding an antioxidant to the medium mimics the effects of *Ct* on *At*. Future experiments could explore the role of different types of ROS and different knockout mutants of *Ct* on the interaction between these two species.

Our second main message is that even in very simple systems like the one we have studied here, several layers of interactions can play out simultaneously. Here we find two layers: competition for the resource, linoleic acid, and facilitation through the removal of the toxin, ROS. Competition is present at all LA concentrations; if we remove environmental toxicity exogenously at high LA, the underlying resource competition is revealed. What we measure under given environmental conditions, then, is the net effect of species on each other, taking into account all the chemical substrates they may be competing for, feeding to one another or removing to facilitate each other’s growth. Because ROS generation is proportional to LA concentration, modifying the initial concentration of just that one compound alters the balance between resource abundance and toxicity, and thereby the dominance of competition or facilitation. This mechanism supports the intuition of the SGH (Fig. 1 in (56)) that these interactions really are context-dependent and do not simply emerge from confounding effects or methodological biases (15). In this system, we have uncovered two mediators or environmental factors (LA and ROS), however it’s possible that there may be more that were not salient enough to be observed. More generally, microbes can construct a surprising number of new niches that could potentially mediate interactions with others (57, 58).

Another important biological question is whether the underlying interactions can help to predict long-term coexistence.

According to early theoretical work, coexistence on a single resource is only possible in small parameter regions in fluctuating environments, when metabolic trade-offs exist (59). Intuitively, therefore, we did not expect coexistence in the competitive scenario at low LA concentration, but over the short time-scale of 5 transfers, our model predicts coexistence, which we recapitulate with the experiment.

It was less clear whether positively interacting species would coexist. Species that interact positively through cross-feeding are expected to coexist if the positive feedback is not too strong and leading to chaotic dynamics (24, 30), but according to our model, two species that are competing for a single resource but facilitating each other through detoxification should not. Eventually, the stronger competitor should dominate. Nevertheless, we show that facilitation by detoxification can increase the duration of coexistence – again in line with the SGH – which may allow the weaker species to survive a limited period of harsh conditions (60, 61). It has been argued that the “expected time to extinction” is an appropriate measure of fitness (62). Nevertheless, according to the model, in longer co-culture experiments, *At* should go extinct at all concentrations (Fig. 4C), which would be an interesting hypothesis to test. One reason why it may not hold and the two species would continue to coexist in the long term, is if there are further interaction layers between the two species like cross-feeding that were not observed in this study.

This brings us to an important discussion point: how much detail is needed to construct a predictive model of a community like the one we have studied here? While we cannot give a definitive answer to this question, our study shows that details of the model matter. First, a consumer-resource (CR) model was needed to capture the context-dependency of interactions, compared to implicitly defining interactions as in a Lotka-Volterra framework (11, 63). Within the CR framework, we found that whether we model a single compound whose effect changes or two compounds – one resource and one toxin – has a huge influence on coexistence (Fig. 4A). More generally, more environmental factors favor coexistence (28). On the one hand, this may be seen as bad news on fitting models to data: without a mechanism for toxicity, it will be difficult to build a good predictive model. On the other hand, some underlying knowledge on the chemistry of the growth medium can go a long way. Does toxicity arise abiotically? Is it dependent on metabolic activity? Distinguishing the two scenarios can be achieved by inoculating bacteria at different time-points into the cell-free medium or into a growing bacterial culture, for example. As long as parameterization is possible, the advantage of such a mechanistic understanding is that it can generate more realistic hypotheses on the addition of further species or changes in resource concentrations, compared to an uninformed or abstract exploration of the parameter space of a model. Making sure enough interaction layers are included in the model to describe the observed experimental pattern requires a tight back and forth between models and data.

As we and others have shown, chemical compounds mediate positive or negative interactions between species, such that even in simple laboratory microcosms, several interaction layers may emerge whose balance and net effect will depend on initial conditions and how the chemical environment changes over time. Whether or not species will coexist in the long-term will not only depend on the sign of these interactions but also the mechanism behind them: are they removing stress for one another or feeding one another? Further verifying the model in this study and understanding how the layering of interactions plays out in the long term are next important steps.

## Methods

### Bacterial species and growth conditions

The species used in this study were *Agrobacterium tumefaciens* str. MWF001 (*At*) and *Comamonas testosteroni* str. MWF001 (*Ct*). More details on these strains can be found in (36). We prepared separate overnight cultures in Tryptic Soy Broth (TSB) starting from a single colony for each species. Cultures were incubated at 28^*°*^C, shaken at 200 rpm. The day after, for each species, we measured the OD_600_ (Ultrospec 10 cell density meter, Amersham Biosciences) and adjusted it to an OD_600_ of 0.05 in 20 ml of fresh TSB, which we incubated for three hours at the same conditions to reach exponential phase. We then measured the OD_600_ to dilute cells to an OD_600_ of 0.1 in 10 ml of minimal medium (MM, Table 1) in 15 ml tubes. We centrifuged these cell suspensions for 20 min at 4’000 rpm at 22^*°*^C, discarded the supernatants and resuspended the pellets in 10 ml of PBS to remove any leftover TSB. We repeated this wash twice, resuspending the pellets in 10 ml of MM. Separately, we prepared 4 ml of the growth media in glass growth tubes, with three replicates per condition for every selected compound as indicated in Table 2. We aliquoted 40 *µ*l of each bacterial culture (from the 10 ml suspended in MM) into the appropriate growth tube containing 4 ml of media to dilute bacteria at 10^5^-10^6^ starting CFU/ml. Growth tubes were then incubated at 28^*°*^C, 200 rpm for 8 days.

**Table 1.**
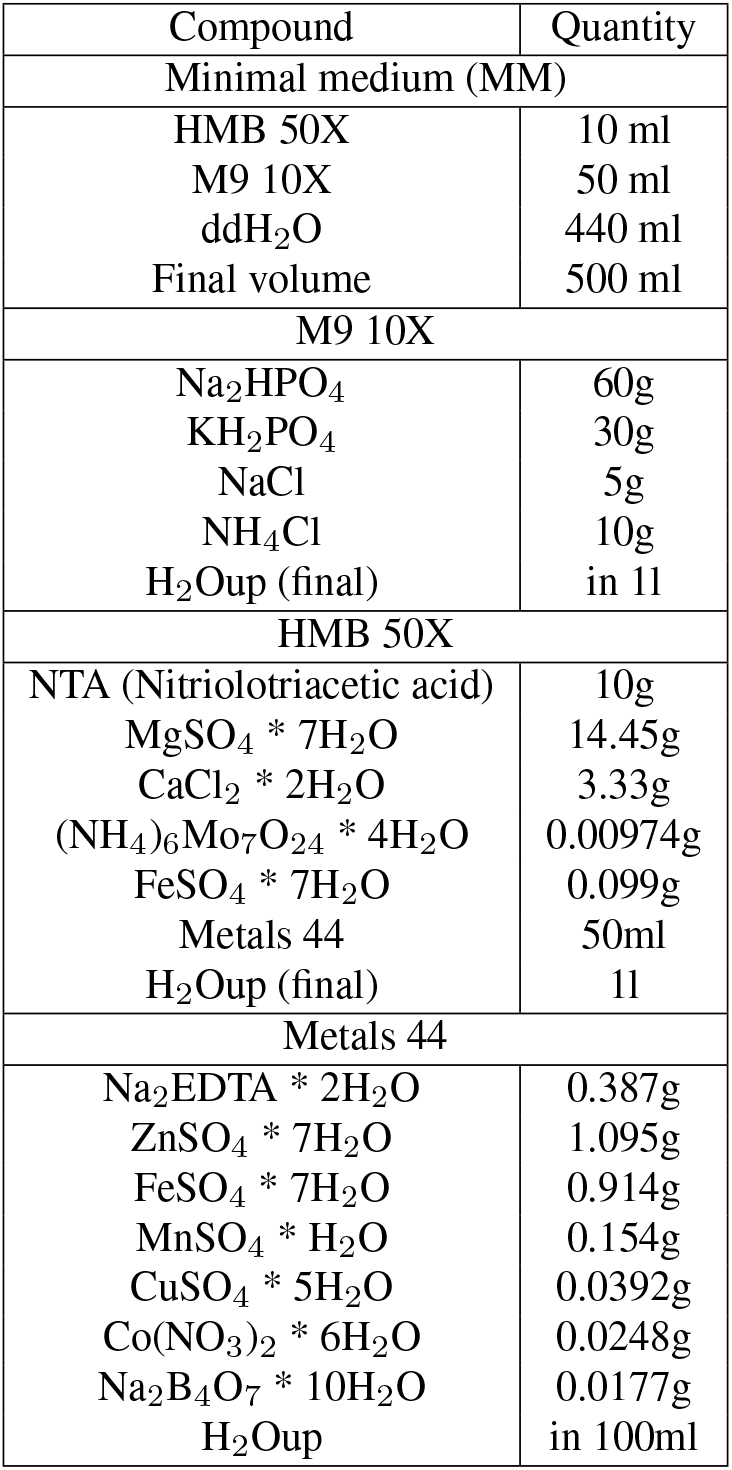
Preparation of minimal medium (MM). The components listed in the table were mixed to prepare stock solutions. The stock solutions were combined according to the protocol to have the final minimal medium (MM) at the top part of the table. H_2_Oup: “filled up to volume”.

| Compound | Quantity |
| --- | --- |
| Minimal medium (MM) |  |
| HMB 50X | 10 ml |
| M9 10X | 50 ml |
| ddH<sub>2</sub>O | 440 ml |
| Final volume | 500 ml |
| M9 10X |  |
| Na<sub>2</sub>HPO<sub>4</sub> | 60g |
| KH<sub>2</sub>PO<sub>4</sub> | 30g |
| NaCl | 5g |
| NH<sub>4</sub>Cl | 10g |
| H<sub>2</sub>Oup (final) | in 1l |
| HMB 50X |  |
| NTA (Nitrilotriacetic acid) | 10g |
| MgSO<sub>4</sub> * 7H<sub>2</sub>O | 14.45g |
| CaCl<sub>2</sub> * 2H<sub>2</sub>O | 3.33g |
| (NH<sub>4</sub>)<sub>6</sub>Mo<sub>7</sub>O<sub>24</sub> * 4H<sub>2</sub>O | 0.00974g |
| FeSO<sub>4</sub> * 7H<sub>2</sub>O | 0.099g |
| Metals 44 | 50ml |
| H<sub>2</sub>Oup (final) | 1l |
| Metals 44 |  |
| Na<sub>2</sub>EDTA * 2H<sub>2</sub>O | 0.387g |
| ZnSO<sub>4</sub> * 7H<sub>2</sub>O | 1.095g |
| FeSO<sub>4</sub> * 7H<sub>2</sub>O | 0.914g |
| MnSO<sub>4</sub> * H<sub>2</sub>O | 0.154g |
| CuSO<sub>4</sub> * 5H<sub>2</sub>O | 0.0392g |
| Co(NO<sub>3</sub>)<sub>2</sub> * 6H<sub>2</sub>O | 0.0248g |
| Na<sub>2</sub>B<sub>4</sub>O<sub>7</sub> * 10H<sub>2</sub>O | 0.0177g |
| H<sub>2</sub>Oup | in 100ml |

**Table 2.**
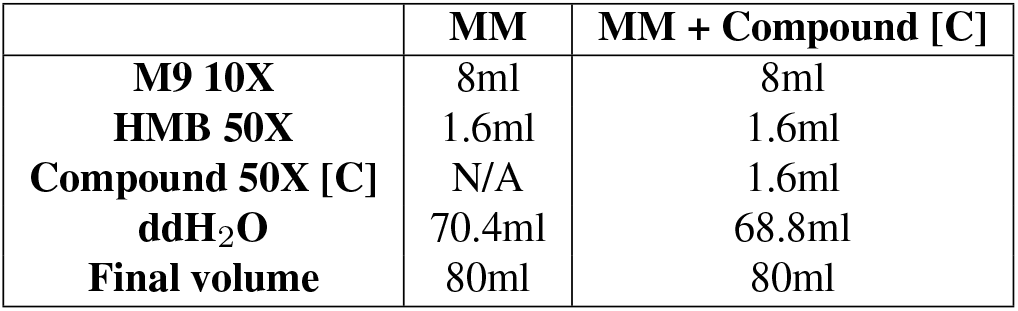
Four different compound concentrations [C] plus a carbon-free medium were prepared by adding the compounds to MM (Table 1). When possible, we prepared 50X concentrated stocks of the tested compounds to have a more standard procedure.

|  | MM | MM + Compound [C] |
| --- | --- | --- |
| **M9 10X** | 8ml | 8ml |
| **HMB 50X** | 1.6ml | 1.6ml |
| **Compound 50X [C]** | N/A | 1.6ml |
| **ddH<sub>2</sub>O** | 70.4ml | 68.8ml |
| **Final volume** | 80ml | 80ml |

### Media preparation and compound storage

The compounds we used for media preparations were: linoleic acid, oleic acid, citric acid, monoethanolamine, petroleum sulfonate, triethanolamine, morpholine, and naphthenic petroleum oil. Naphthenic petroleum oil and petroleum sulfonate were kindly given to us by Peter Kueenzi at Blaser Swisslube. All compounds were stored according to indications provided by the manufacturer: Linoleic and oleic acid were stored in individual 1 ml aliquots at -20^*°*^C. Each aliquot was single-used to avoid multiple thaw-freeze processes, which could affect compound stability. We discarded and replaced aliquots that were more than 1 year old.

All growth media were prepared using a carbon-free minimal medium (MM, Table 1), to which we added the appropriate concentrations of different compounds (or none) (Table 2). When possible, we prepared 50-fold concentrated stocks in water to standardize media preparation. Linoleic acid, oleic acid, petroleum sulfonate and naphthenic petroleum oil are not fully miscible in water, and we used them as constantly shaken emulsions. This also prevented the preparation of stock solutions, and we therefore aliquoted the adequate amounts directly from their 99% pure stock (ddH_2_O concentrations were adjusted accordingly as in Table 2). For linoleic acid and oleic acid, we thawed aliquots at room temperature before proceeding with media preparation. All assembled compound-supplemented media were incubated at 28^*°*^C, 200 rpm for 3 hours to allow complete mixing before distributing them into glass growth tubes and inoculating them with bacteria as described above.

### Transfer experiments

We grew both *At* and *Ct* mono- and co-cultures for 72 hours in both 0.1% LA and 0.75% LA. After 72 hours, we transferred a 1% aliquot of each culture (40 *µ*l) into 4 ml of fresh medium and grew bacteria for another cycle of 72 hours. We performed five transfers, quantifying population sizes at each transfer as described next.

### Quantification of population size

To quantify bacterial population sizes over time, we took 20 *µ*l aliquots from each growth tube, performed serial dilutions in 96-well plates filled with 180 *µ*l of PBS and plated them on tryptic soy agar (TSA) plates or on lysogeny broth (LB) agar (depending on availability, no differences were observed). Plates were incubated at 28^*°*^C. *Ct* and *At* formed countable colonies (CFUs) after 24 or 48 hours of incubation, respectively. To distinguish *At* and *Ct* when growing in the co-culture, bacteria were also plated on LB agar supplemented with 14.25*µ*g/ml of sulfamethoxazole and 0.75*µ*g/ml of trimethoprim to count only *At* colonies. Moreover, the GFP marker of *At* further helped to differentiate *At* and *Ct* colonies.

### Quantification of compound effect on bacteria

CFU counts were used to plot growth curves of CFU/ml over time and we calculated the area under the growth curve (AUC) to have a better comparison between the different tested conditions and their MM control. We calculated the ratio between the AUC of each replicate per condition and the mean of the AUC of the three MM control replicates, and used the log_2_- fold change of these data to build heatmaps showing the effect of each compound on each of the species (Fig. 1). Raw data for all growth curves is shown in Fig. S2. T-tests were performed to compare the tested conditions to the MM control.

### ROS detection assay

We used the Thiobarbituric Acid Reactive Substances (TBARS) assay to indirectly assess the presence of ROS-induced oxidative stress as described in ref. (46). Malondialdehyde (MDA) is the primary product of lipid peroxidation, the oxidative degradation induced by ROS. If there is ROS-induced degradation of LA in our media, this process would lead to MDA production. The TBARS assay measures the formation of the new adduct MDA-TBA2 upon reaction between the MDA in the medium and supplemented thiobarbituric acid (TBA). MDA-TBA2 presence is measured by its absorbance at 532 nm and the detected values are transformed in MDA-TBA2 concentrations through interpolation with a calibration curve built using eight MDA-TBA2 standards at known concentrations (Fig. S1). All reacting solutions necessary for this assay were prepared following the detailed protocol in (46).

### Mathematical model

#### Equations and fitting

We fitted the data from the experiments to mathematical models to determine whether species are expected to compete, facilitate each other, and whether they would coexist over long term serial transfers. Details are given in supplementary material.

In the first model, bacterial species abundance *B* over time depends on the concentration of LA *C* according to its consumption through a Monod uptake function, with maximum growth rate *r*, half-saturation constant *K*, and yield *Y*. LA also induces mortality depending on its concentration. We assumed that its toxicity increases linearly over time and is proportional to the LA concentration, leading to the linear expression (*β* + *γt*)*C*. This linear increase may appear arbitrary, but it was needed to get a reasonable fit to the monoculture of *At* at 0.75% LA, where the population grows initially and then drops abruptly. LA concentration *C* varies only due to consumption by bacteria. This formulation is similar to the classic growth-inhibiting substrate approach as described in example 1 from (24) with a hump-shaped functional response (Haldane or Type IV), except that it allows for a negative growth rate (death) while the Haldane form tends to 0. The equations for the variation of bacterial abundance and LA concentration in a monoculture are:

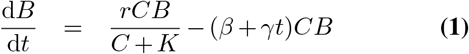

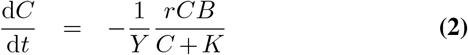

The equations for two species *B*_1_ and *B*_2_ in co-culture in LA are given in the supplementary material. We used this model to fit the growth of *At* and *Ct* in monoculture in a range of concentrations of LA (0.05%, 0.1%, 0.5% and 0.075%) and compared the predictions for the abundances of the two species to the experimental data in a short-term co-culture and their long-term coexistence over transfers. In this first model, the estimated parameters were *r*_*C*1_, *r*_*C*2_, *Y*_*C*1_, *Y*_*C*2_, *K*_*C*1_, *K*_*C*1_ and the toxicity parameters for *At β*_2_ and *γ*_2_, as we set *β*_1_ and *γ*_1_ to zero for *Ct*. The best-fit estimates are listed in supplementary material.

We then used a second model that accounts for the production of ROS by LA oxidation. The equations for the monoculture growth are now:

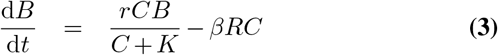

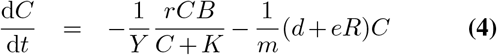

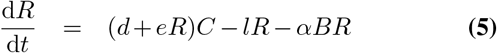

Because we had acquired data on the spontaneous oxidation of LA in cell-free media, we could first estimate the parameters *d, e, m*, and *l* using a ROS proxy measured at different LA concentrations. We then fixed these parameters as estimated from the monoculture and further estimated the parameters of growth, toxicity, and detoxification for each species. All parameter estimations were obtained using the modFit function from FME package (version 1.3.6.1) in R version 4.1.0.

#### Comparing model predictions to co-cultures and transfers

For both models, we used the parameters obtained from the estimation of the monoculture data to predict co-culture dynamics, and compared them to the actual co-culture data, using Eq. (S9). We also simulated the serial transfers with varying dilution rates and initial LA concentrations to predict the likelihood of coexistence of the two species over time. The transfer parameter sweeps were coded in C++. In the second model, we mimicked the addition of a ROS quencher to the media by setting initial ROS concentration to zero, as well as parameters *d, e*, and *l* and compared the predicted dynamics to the actual data using TBHQ.

## ACKNOWLEDGEMENTS

We thank Andrew Quinn and Philipp Engel for their help to obtain metabolomics data that is not included in the manuscript, and Katia Annen for help with the experiments. We thank Peter Kueenzi at Blaser Swisslube for the chemical compounds used to generate the data in Figure 1. We thank Afra Salazar de Dios, Julien Luneau, Margaret Vogel, Oliver Meacock and Massimo Amicone for insightful comments on the manuscript..

