## Supplementary notes, figures and table for "Oxidative stress changes interactions between two bacterial species"

### Supplementary Note A: Model details and fitting

**Model with implicit toxicity (Model 1).** We fit the data from the experiments to a mathematical model to predict whether species are expected to engage in competition, facilitation, and/or to coexist over long term serial transfers.

The first model is a modified Monod model with maximum growth rate $r$, half-saturation constant $K$, and yield $Y$, in which we incorporate a mortality term to take into account concentration-dependent toxicity of linoleic acid (LA). We assume that toxicity of the environment $T(t)$ increases linearly over time and is proportional to the LA concentration $T(t) = (\beta + \gamma t)$.

The simplest equations that we start from are for a single species *B* in a batch culture with LA (*C*):

$$\frac{\mathrm{d}B}{\mathrm{d}t} = \left(\frac{r}{C(t)+K} - (\beta + \gamma t)\right) C(t)B(t) \quad \textbf{(S1)}$$

$$\frac{\mathrm{d}C}{\mathrm{d}t} = -\frac{1}{Y}\frac{r}{C(t)+K}C(t)B(t) \quad \textbf{(S2)}$$

The equations for two species ($B_1$, corresponding to *Ct* and $B_2$, corresponding to *At*) in coculture in LA are:

$$\frac{\mathrm{d}B_1}{\mathrm{d}t} = \left(\frac{r_1}{C(t)+K_1} - (\beta_1 + \gamma_1 t)\right) C(t)B_1(t) \quad \textbf{(S3)}$$

$$\frac{\mathrm{d}B_2}{\mathrm{d}t} = \left(\frac{r_2}{C(t)+K_2} - (\beta_2 + \gamma_2 t)\right) C(t)B_2(t) \quad \textbf{(S4)}$$

$$\frac{\mathrm{d}C}{\mathrm{d}t} = -\frac{1}{Y_1}\frac{r_1}{C(t)+K_1}C(t)B_1(t) - \frac{1}{Y_2}\frac{r_2}{C(t)+K_2}C(t)B_2(t) \quad \textbf{(S5)}$$

Because the bacteria show some growth in the minimal medium, we assume an additional unknown nutrient to be present in the minimal medium, whose concentration $N(t)$ is modelled in an additional equation. The updated model becomes:

$$\frac{\mathrm{d}B_1}{\mathrm{d}t} = \left(\frac{r_{C1}}{C(t)+K_{C1}} - (\beta_1 + \gamma_1 t)\right) C(t)B_1(t) + \frac{r_{N1}}{N(t)+K_{N1}}N(t)B_1(t) \quad \textbf{(S6)}$$

$$\frac{\mathrm{d}B_2}{\mathrm{d}t} = \left(\frac{r_{C2}}{C(t)+K_{C2}} - (\beta_2 + \gamma_2 t)\right) C(t)B_2(t) + \frac{r_{N2}}{N(t)+K_{N2}}N(t)B_2(t) \quad \textbf{(S7)}$$

$$\frac{\mathrm{d}C}{\mathrm{d}t} = -\frac{1}{Y_{C1}}\frac{r_{C1}}{C(t)+K_{C1}}C(t)B_1(t) - \frac{1}{Y_{C2}}\frac{r_{C2}}{C(t)+K_{C2}}C(t)B_2(t) \quad \textbf{(S8)}$$

$$\frac{\mathrm{d}N}{\mathrm{d}t} = -\frac{1}{Y_{N1}}\frac{r_N 1}{C(t)+K_{N1}}C(t)B_1(t) - \frac{1}{Y_{N2}}\frac{r_{N2}}{C(t)+K_{N2}}C(t)B_2(t) \quad \textbf{(S9)}$$

We then first estimate the parameters of the growth in the minimal medium using an arbitrary concentration for this unknown nutrient (0.01), by using the data from both mono- and cocultures of *Ct* (species $B_1$) and *At* (species $B_2$) in the minimal medium. This allows us to obtain estimates for the parameters $r_{N1}$, $r_{N2}$, $Y_{N1}$, $Y_{N2}$, $K_{N1}$ and $K_{N1}$. Then, we fix these parameters and estimate the parameters for the growth of *At* and *Ct* in monoculture using a range of concentrations of LA (0.05%, 0.1%, 0.5% and 0.075%). This allows us to estimate $r_{C1}$, $r_{C2}$, $Y_{C1}$, $Y_{C2}$, $K_{C1}$, $K_{C1}$ and the toxicity parameters for *Ct* ($\beta_1$ and $\gamma_1$) and *At* ($\beta_2$ and $\gamma_2$). .

The best-fit parameters estimates obtained from the modFit run for *Ct* with model 1 are $r_{C1} = 2.378$, $K_{C1} = 0.0006$, $Y_{C1} = 1.383e9$, $\beta_1 = 0.299$, $\gamma_1 = 3.014e-6$ and minimal medium parameters are $r_{N1} = 346.5$, $K_{N1} = 0.101$, $Y_{N1} = 1.278e9$.

The parameters for *At* are: $r_{C2} = 1.941$, $K_{C2} = 0.001$, $Y_{C2} = 2.383e9$, $\beta_2 = 3.359$, $\gamma_2 = 0.563$ and minimal medium parameters are $r_{N2} = 891.4$, $K_{N2} = 1.013$, $Y_{N2} = 3.139e8$.

**Model with explicit toxicity due to ROS (Model 2).** In this second model, we add a new state variable corresponding to the concentration of ROS. LA is now only a nutrient, and the toxicity is proportional to ROS concentration (which can increase), so we do not need a specific parameter for the toxicity accumulation. The parameters $\beta_1$ and $\beta_2$ are the sensitivity of *Ct* and *At* to ROS (high value meaning low tolerance). The uptake of LA does not change from the previous model. The ROS intrinsic dynamics depend on their production by the oxidation of LA (spontaneous oxidation at rate $d$, positive feedback by ROS presence in the media, $e$, their half-life $l$ and the yield of ROS production $m$). To this intrinsic part, we add the detoxification by the cells, through parameters $\alpha_1$ for *Ct* and $\alpha_2$ for *At*.

DRAFT

The coculture equations become:

$$\frac{\mathrm{d}B_1}{\mathrm{d}t} = \frac{r_{C1}}{C(t)+K_{C1}}C(t)B_1(t) - \beta_1 B_1(t)R(t) + \frac{r_{N1}}{N(t)+K_{N1}}N(t)B_1(t) \tag{S10}$$

$$\frac{\mathrm{d}B_2}{\mathrm{d}t} = \frac{r_{C2}}{C(t)+K_{C2}}C(t)B_2(t) - \beta_2 B_2(t)R(t) + \frac{r_{N2}}{N(t)+K_{N2}}N(t)B_2(t) \tag{S11}$$

$$\frac{\mathrm{d}C}{\mathrm{d}t} = -\frac{1}{Y_{C1}}\frac{r_{C1}}{C(t)+K_{C1}}C(t)B_1(t) - \frac{1}{Y_{C2}}\frac{r_{C2}}{C(t)+K_{C2}}C(t)B_2(t) - \frac{1}{m}(d+eR(t))C(t) \tag{S12}$$

$$\frac{\mathrm{d}N}{\mathrm{d}t} = -\frac{1}{Y_{N1}}\frac{r_N 1}{C(t)+K_{N1}}C(t)B_1(t) - \frac{1}{Y_{N2}}\frac{r_{N2}}{C(t)+K_{N2}}C(t)B_2(t) \tag{S13}$$

$$\frac{\mathrm{d}R}{\mathrm{d}t} = (d+eR(t))C(t) - lR(t) - \alpha_1 B_1(t)R(t) - \alpha_2 B_2(t)R(t) \tag{S14}$$

The monoculture equations can be derived by setting one of the two bacterial densities to zero. Because we now have data on the spontaneous oxidation of LA in cell-free media, we can first estimate the parameters $d$, $e$, $m$, and $l$ using the ROS proxy at different LA concentrations. We then use these fixed parameters to estimate the parameters of growth, toxicity, and detoxification for single species in monoculture. Then, predictions can be made, as in the previous model, for the co-culture dynamics, short- and long-term dynamics (serial transfers), and mimicking the addition of a ROS quencher to the media (setting initial ROS concentration to zero, as well as parameters $d$, $e$, and $l$).
The best-fit parameter estimates for ROS intrinsic dynamics in model 2 are: $m = 0.879$, $d = 0.114$, $e = 1.94$, $l = 0.346$. Parameters for *Ct* are: $r_{N1} = 5.518$, $K_{N1} = 5.518$, $Y_{N1} = 4.08e8$, $\beta_1 = 8.58$, $\alpha_1 = 4.61e-6$. Parameters for *At* are: $r_{N2} = 126.5$, $K_{N2} = 0.357$, $Y_{N2} = 3.27e8$, $\beta_2 = 20$, $\alpha_2 = 0$.

DRAFT

### Supplementary Note B: Supplementary figures

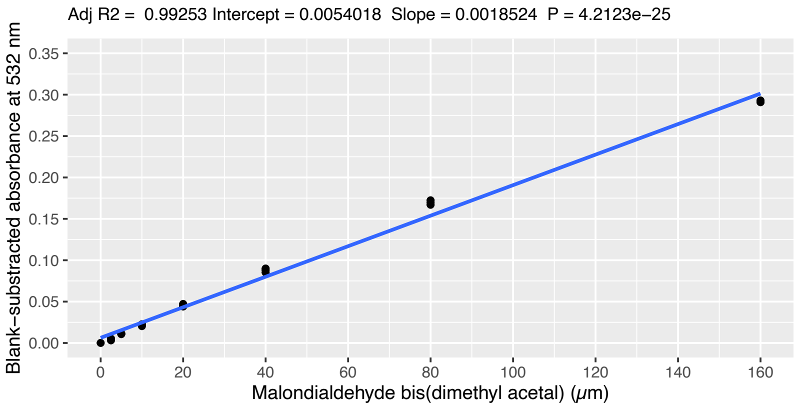


**Fig. S1.** Calibration curve ROS quantification. Malondialdehyde bis(dimethyl acetal) (MDA) concentration is used as a proxy for ROS accumulation: the higher the MDA concentration, the higher is the ROS abundance in the sample (46). We chose six increasing concentrations of MDA (0, 2.5, 5, 10, 20, 40, 80, 160 µM) and we used three replicates per concentration to build a calibration curve. We performed the TBARS assay on each calibration sample as described in (46) and measured the 532nm absorbance. We subtracted the absorbance of the blank (0 µM MDA) from each calibration sample and plotted MDA concentration versus blank-subtracted absorbance. We obtained a calibration curve and used its parameters to calculate the MDA concentration of the experimental samples shown in Fig. 3.

| Enzyme family | Gene names | Annotated genes in *At* str. MWF001 | Annotated genes in *Ct* str. MWF001 |
|---|---|---|---|
| Superoxide dismutases | SodA, SodM | SodB (3) | SodB (2) |
| Catalases | KatA, KatE, KatG | KatG (1) | KatG (1) |
| Thiol peroxidase, thioredoxins | TpxD, TlpA, Etrx, TrxA, TrxB | TlpA (1), TrxA, TrxB (2), TrxC | TrxA (4), TrxC |
| Peroxiredoxin | AhpC, AhpD, Bcp | AhpC, AhpD, Bcp (2) | AhpC, AhpF, BcpB, Bcp |
| Glutathione reductase, glutaredoxin | gor, grxA | grxC, grxD | grxC, grxD |

**Table S1.** ROS resistance genes in *At* and *Ct*. We searched for a list of putative ROS-degrading enzymes from a recent review paper (55) by searching the annotated genes of our two strains. We show the gene families listed in (55), and whether we found genes of the same family in our two genomes with gene number shown in brackets. This analysis shows that gene presence/absence tells us little about which of the two strains is more resistant to ROS.

DRAFT

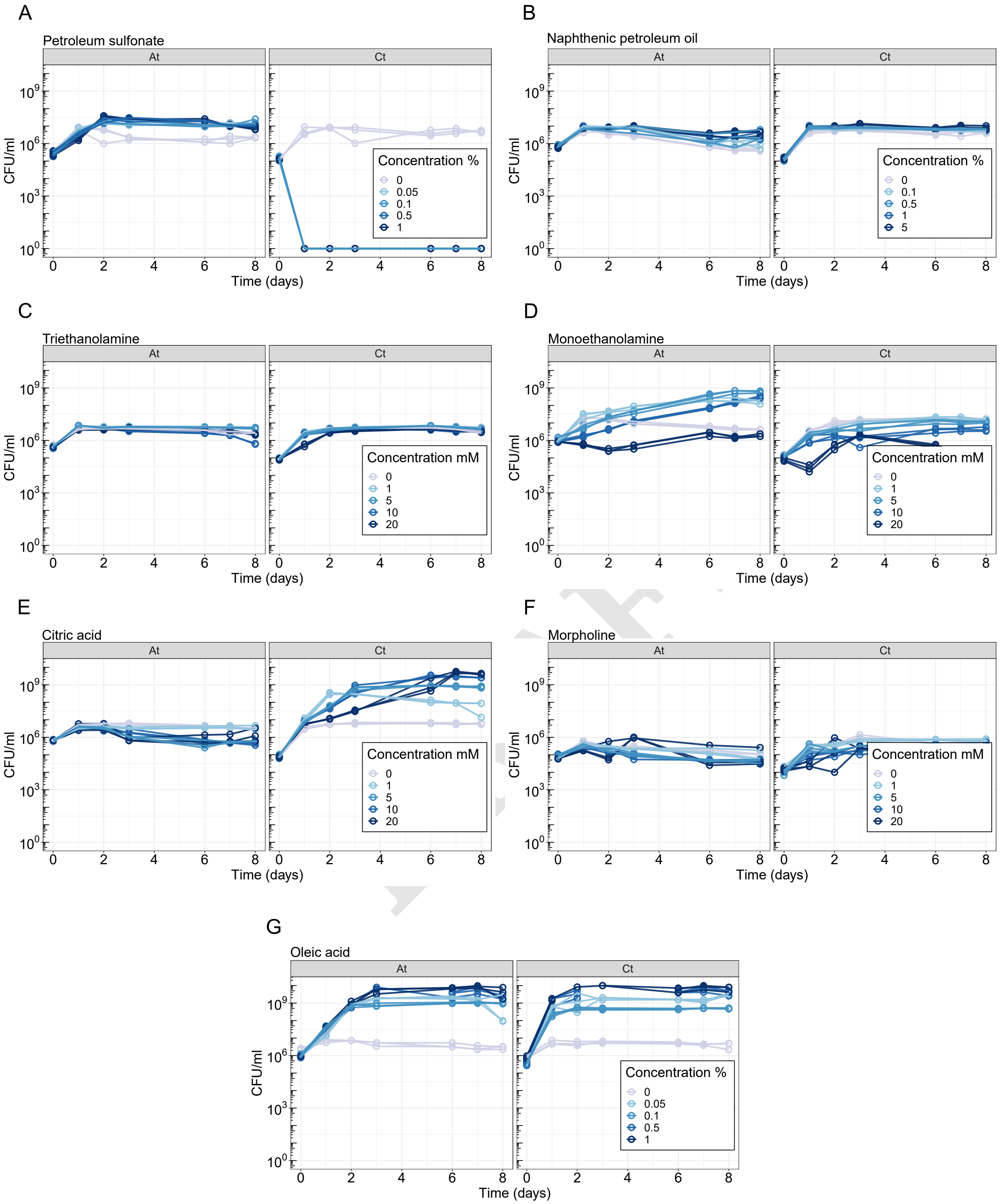


**Fig. S2.** Growth curves of *At* and *Ct* in MWF compounds. We chose compounds representative of MWF composition and grew *At* and *Ct* in increasing concentrations of the following compounds: petroleum sulfonate (A), naphthenic petroleum oil (B), triethanolamine (C), monoethanolamine (D), citric acid (E), morpholine (F) and oleic acid (G). Darker gradient of blue curves represents increasing compound concentration.
